# Dynamic Brain Connectivity Alternation Detection via Matrix-variate Differential Network Model

**DOI:** 10.1101/446237

**Authors:** Jiadong Ji, Yong He, Lei Xie

## Abstract

**Motivation:** Nowadays brain connectivity analysis has attracted tremendous attention and has been at the foreground of neuroscience research. Brain functional connectivity reveals the synchronization of brain systems through correlations in neurophysiological measures of brain activity. Growing evidence now suggests that the brain connectivity network experiences alternations with the presence of numerous neurological disorders, thus differential brain network analysis may provides new insights into disease pathologies. For the matrix-valued data in brain connectivity analysis, existing graphical model estimation methods assume a vector normal distribution that in essence requires the columns of the matrix data to be independent. It is obviously not true, they have limited applications. Among the few solutions on graphical model estimation under a matrix normal distribution, none of them tackle the estimation of differential graphs across different populations. This motivates us to consider the differential network for matrix-variate data to detect the brain connectivity alternation.

**Results:** The primary interest is to detect spatial locations where the connectivity, in terms of the spatial partial correlation, differ across the two groups. To detect the brain connectivity alternation, we innovatively propose a Matrix-Variate Differential Network (MVDN) model. MVDN assumes that the matrix-variate data follows a matrix-normal distribution. We exploit the D-trace loss function and a Lasso-type penalty to directly estimate the spatial differential partial correlation matrix where the temporal information is fully excavated. We propose an ADMM algorithm for the Lasso penalized D-trace loss optimization problem. We investigate theoretical properties of the estimator. We show that under mild and regular conditions, the proposed method can identify all differential edges accurately with probability tending to 1 in high-dimensional setting where dimensions of matrix-valued data *p, q* and sample size *n* are all allowed to go to infinity. Simulation studies demonstrate that MVDN provides more accurate differential network estimation than that achieved by other state-of-the-art methods. We apply MVDN to Electroencephalography (EEG) dataset, which consists of 77 alcoholic individuals and 45 controls. The hub genes and differential interaction patterns identified are consistent with existing experimental studies.

**Supplementary information:** Supplementary data are available online.

## 1 Introduction

Nowadays brain connectivity analysis has attracted tremendous attention and has been at the foreground of neuroscience research. Brain functional connectivity reveals the synchronization of brain systems through correlations in neurophysiological measures of brain activity. Varoquaux and Craddock (2013) pointed out that when measured during the resting state, it maps the intrinsic functional architecture of the brain. Growing evidence now suggests that the brain connectivity network experiences alternations with the presence of numerous neurological disorders, including Alzheimer’s disease, attention deficit hyperactivity disorder, autism spectrum disorder (Fox and Greicius, 2010; Daliri and Behroozi, 2013). In this paper, we focus on the problem of comparing brain functional connectivity patterns across the diseased versus the healthy control, which helps to provide new insights into disease pathologies.

Graphical model has been widely adopted to capture the interactions (connectivity) between variables, especially the Gaussian Graphical Model (GGM). Under GGM, two variables interact if and only if the corresponding entry of the precision matrix is nonzero, where the precision matrix is defined as the inverse of the covariance matrix. In recent years, numerous research on precision matrix estimation has appeared in the high dimensional setting, for example, Meinshausen and Bühlmann (2006); Yuan and Lin (2007); Cai *et al.* (2011); Zhang and Zou (2014). Some researchers studied a more flexible Gaussian copula graphical model which relax the restrictive Gaussian assumption of GGM in real application, for instance, Liu *et al.* (2009, 2012); He *et al.* (2017a). This research area is very active, as a result, this list of references here is illustrative rather than comprehensive.

In the brain connectivity study, it is often the case that the functional MRI data are collected for multiple subjects from the disease group and normal control. Compared to focusing on the brain region network from one particular group, it is of greater interest to investigate how the network of connected brain regions alters from disease group to the control group. It provides deeper insights on an underlying biological process. In other word, it is more appropriate to model the differential network of brain regions between two groups. Recently, differential network modelling has emerged as an important tool to analyze a set of changes in graph structure between two groups, and is typically modelled as the difference of the precision matrices for each group, for example, Zhao *et al.* (2014); Xia *et al.* (2015); Yuan *et al.* (2015) and so on. All these works assume that the vector of interacting variables follows a normal distribution. However, the observed functional MRI data in brain functional connectivity study is in the matrix form, whose row variables represents the brain regions while the column variables represents time points. Typically, the number of brain regions is of the order 10^2^ and the number of time points is around 150 to 200. In fact, the matrix-valued data have been emerging rapidly in the current big data era. To our knowledge, for the type of matrix-variate observation data, there has not been any work on analyzing the differential network, which motivates us to propose a corresponding differential network framework.

For the matrix-valued data, matrix normal distribution is becoming increasingly popular in modelling such observations. In the matrix normal distribution framework, the brain connectivity network analysis can be viewed as a precision matrix inference problem. Specifically, let ***X***_*p* × *q*_ be the spatial-temporal matrix data from an image modality. We say***X***_*p* × *q*_ follows a matrix normal distribution with the Kronecker product covariance structure **Σ** = **Σ**_*T*_ ⊗ **Σ**_*S*_, which we write as

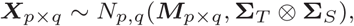

if and only if Vec(***X***_*p* × *q*_) follows a multivariate normal distribution with mean Vec(***M***_*p* × *q*_) and covariance **Σ** = **Σ**_*T*_ ⊕ **Σ**_*S*_, where Vec(***X***_*p* × *q*_) is formed by stacking the columns of Vec(***X***_*p* × *q*_) into a vector in ℝ^*pq*^, **Σ**_*S*_ ∈ℝ^*p* × *p*^ and **Σ**_*T*_ ∈ ℝ^*q × q*^ denote the covariance matrices of *p* spatial locations and *q* times points, respectively. Correspondingly, we have

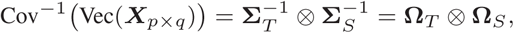

where **Ω**_*S*_ ∈ ℝ^*p × p*^ and **Ω**_*T*_ ∈ ℝ^*q × q*^ denote the spatial and temporal precision matrix, respectively. Note that **Σ**_*S*_ and **Σ**_*T*_ are only identifiable up to a scaled factor, as 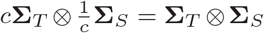 for any *c* > 0. In fact, in the brain connected network analysis, the partial correlation is a commonly adopted correlation measure (Peng *et al.*, 2009; Zhu and Li, 2018). Besides, the primary interest in brain connectivity analysis is to infer the the connectivity network characterized by the spatial precision matrix **Ω**_*S*_ while the temporal precision matrix **Ω**_*T*_ is of little interest and thus can be viewed as a nuisance parameter. In a word, under the matrix norm framework a region-by-region spatial partial correlation matrix characterizes the brain connectivity graph, in which nodes represent brain regions, and links measure conditional dependence between the brain regions. Brain connectivity analysis is equivalently transformed into the estimation of spatial partial correlation matrix. We remark here that the matrix normal distribution framework has been widely adopted in real application and is scientifically plausible in neuroimaging analysis. (see, for example, Yin and Li, 2012; Leng and Tang, 2012; Zhou, 2014; Xia and Li, 2017; Zhu and Li, 2018; Xia and Li, 2018)

None of existing graphical models under a matrix normal distribution tackle the differential network between the disease group and the control group directly, but instead focus on only a single graph (Zhu and Li (2018) focused on joint estimation of multiple graphs). In this paper, we for the first time propose a differential network model for dynamic matrix-variate data paralleled to static vector-variate data. In specific, assume ***X***_*p* × *q*_ ~ *N_p,q_* (***M***_*p* × *q*_, **Σ***_T_X__* ⊗ **Σ***_S_X__*) be the spatial-temporal matrix data from the disease group and *Y_p × q_* ~ *N_p,q_* (***M***_*p* × *q*_, **Σ***_T_Y__* ⊗ **Σ***_S_Y__*) from the control group. The primary interest in brain connectivity alternation detection is to infer the the differential connectivity network characterized by the difference of spatial precision matrices from two groups, namely, **Ω***_S_X__* — **Ω***_S_Y__*. For the matrix-valued data, directly applying the existing differential network estimation method, which assumes the row variables follow a vector normal distribution, would ignore the dependence between the columns of the matrix data and result in uncorrect conclusion. For example, the columns in functional magnetic resonance imaging (FMRI) study corresponds to times series of repeatedly measured brain activity and are highly correlated. FMRI is one of the mainstream imaging modalities to study brain functional connectivity nowadays. As an alternative, whitening can reduce the between-column correlation. However, Zhu and Li (2018) showed that their joint estimation method substantially outperforms the state of the art vector-normal-based differential network estimation methods such as Zhao *et al.* (2014) and Yuan *et al.* (2015), facilitated by whitening. In the simulation study, we show that the proposed method still outperforms Zhu and Li (2018)’s joint estimation methods, which further indicate that methods facilitated by whitening are inferior to the proposed method.

The main contribution of the current work lies in the following aspects. Firstly, we innovatively propose a Matrix-Variate Differential Network (MVDN) model which is particularly useful in modelling connectivity alternation for matrix-variate data such as in brain connectivity analysis. Secondly, we propose a computationally efficient algorithm for estimating the MVDN, which directly estimate the spatial partial correlation matrix difference without attempting to estimate the partial correlation matrices separately. Thirdly, the proposed approach fully excavates the μtemporal information without assuming the columns of the matrix data to be independent compared with the vector-normal graphical model methods. At last, we show that the proposed method can identify all differential edges accurately with probability tending to 1 under mild and regular conditions as *p, q* and sample size *n* go to infinity.

We denote some notations which are adopted throughout of the paper. For any vector ***μ*** = (*μ*_1_,…, *μ_d_*) ∈ ℝ*^d^*, let 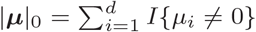, 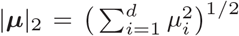 and |***μ***|_∞_ = max_*i*_|*μ_i_*. Let ***A*** = [*a_ij_*] ∈ ℝ^*d × d*^, define 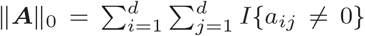 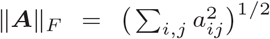, ‖***A***‖_∞_ = max_*i,j*_|a_*ij*_| and ‖***A***‖_1_ = 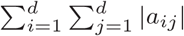. We use λ_min_(***A***) and λ_max_(***A***) to denote the smallest and largest eigenvalues of ***A*** respectively, *κ*(***A***) denote the conditional number of ***A*** and denote the trace of ***A*** as Tr(***A***) and det(***A***) be the determinant of ***A***. The notation ⊗ represents Kronecker product. For a set 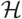, denote by 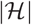 the cardinality of 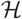. For a real number *x*, denote by ⌊*x*⌋ the largest integer smaller than or equal to *x* and define *p* ∨ *q* = max(*p, q*).

The remainder of the article is organized as follows. In Section 2, we present our matrix-variate differential network modeling framework and the ADMM algorithm for the Lasso penalized D-trace loss optimization problem. We also introduce the simulation designs and real data sets in this section. In section 3, we provide the theoretical results, with the detailed assumption and proofs delegated to the supplementary material. We also present the simulation results and the real data application results in this section. At last, we give a brief discussion on possible future directions in Section 4.

## 2 Methods

Let ***X, Y*** ∈ ℝ^*p* × *q*^ be denote the spatial-temporal matrices of the diseased and the healthy control groups. We assume***X,Y*** follows a matrix normal distribution with the Kronecker product covariance structure, and the mean matrices, without loss of generality, are assumed to be zero. In other words,***X*** ~ *N_p,q_*(**0**_*p×q*_,**Σ**_*T_X_*_ ⊗ **Σ***_S_X__*), and ***Y*** ~ *N_p,q_*(**0**_*p×q*_,**Σ**_*T_Y_*_ ⊗ **Σ***_S_Y__*). Under the matrix normal framework, let

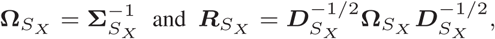

where ***R***_*S_X_*_ is the partial correlation matrix of the spatial locations in diseased group and ***D***_*S_X_*_ is the diagonal matrix of **Ω**_*S_X_*_. Similarly, for the healthy control group, we can define **Ω**_*S_Y_*_ and ***R****_S_Y__* parallelly. The primary interest of this paper is to detect spatial locations where the connectivity, in terms of the spatial partial correlation, differ across the two groups. In this paper, we define the differential network between two groups as the difference between the two spatial partial correlation matrices, denoted by **Δ** = ***R****_S_Y__* — ***R****_S_X__*. The **Σ**_*T_X_*_, **Σ**_*T_Y_*_ are viewed as nuisance parameters.

Let {***X***_1_, …, ***X***_*n*1_} and {***Y***_1_,…, ***Y***_*n*2_}, each a matrix with dimension *p × q*, be two sets of i.i.d. random samples from the independent matrix normal populations ***X*** and ***Y***, respectively. Motivated by the D-trace loss function in Yuan *et al.* (2015), we propose to estimate **Δ** by minimizing the following loss function:

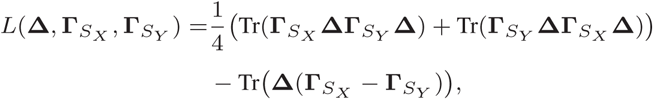

where **Γ**_*S_X_*_ and **Γ**_*S_X_*_ denote the correlation versions of **Σ**_*S_X_*_ and **Σ**_*S_X_*_, respectively. It can be shown that the Hessian matrix with respect to **Δ** of the loss function is (**Γ**_*S_X_*_ ⊗ **Γ**_*S_X_*_ + **Γ**_*S_Y_*_ ⊗ **Γ**_*S_X_*_)/2, which indicates that the loss function *L*(**Δ**, **Γ**_*S_X_*_, **Γ**_*S_Y_*_) is convex with respect to **Δ** and has a unique minimizer at **Δ** = ***R***_*S_Y_*_ — ***R***_*S_X_*_. In brain connectivity alternation detection analysis, it is often the case that the altered connectivity in the spatial networks of two groups is far less compared with the dimensionality, which motivates us to add a penalty to the D-trace loss function. Generally, we can add a decomposable non-convex penalty function which has the form *P*_λ_ (***δ***) = Σ_*j,k*_ *p*_λ_ (Δ_*j,k*_) such as smoothly clipped absolute deviation (SCAD) penalty. For simplicity, we consider a simple lasso penalty here. Finally **Δ** is estimated by solving the following optimization problem:

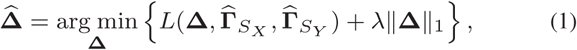

where λ is a tuning parameter and 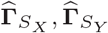 are sample correlation matrices defined as

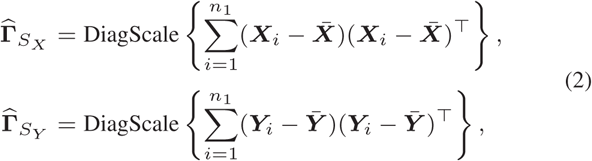

where for a square matrix ***C***, 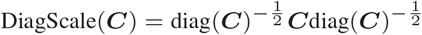. Note that the sample correlation matrices in (2) are based on the matrix-valued observations. The optimization problem in (1) can be solved by the Alternating Direction method of Multipliers (ADMM) algorithm. To present the detailed procedure, we first introduce two multivariate functions *G*(·, ·, ·, ·) and *H*(·, ·). For symmetric *p × p* matrices ***A*** and ***B***, let 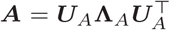 and 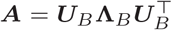 be the corresponding eigenvalue decompositions respectively, where 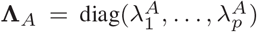 and 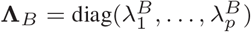. Define function *G*(·, ·, ·, ·) as

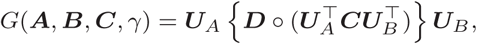

where ***D*** = (*D_ij_*) with 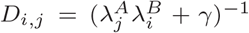 and ○ denotes the Hadamard product of two matrices. For a positive λ, define a ℝ^*p × p*^ · ℝ → ℝ^*p × p*^ map *H*(·, ·) as follows:

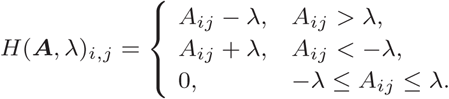

In detail, we summarize the procedure in Algorithm 1.

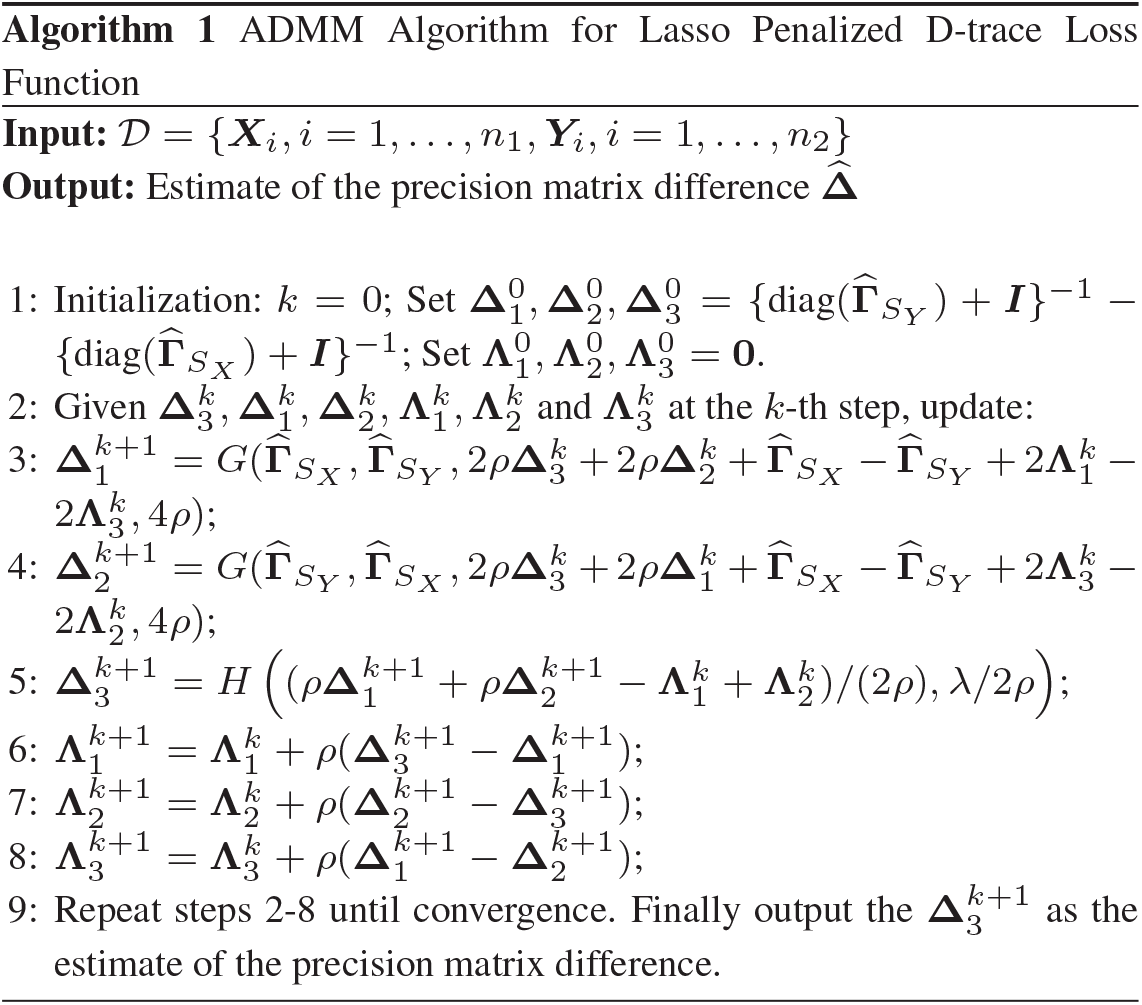

In the simulation study and real data analysis, we set the step size *ρ* = 1 and terminate the algorithm if in the *k* + 1-th step, we have

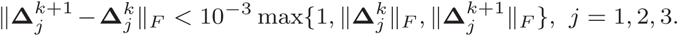

The tuning parameter λ is selected by minimizing the Bayesian information criterion, i.e., λ is chosen to minimize

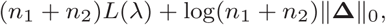

where *L*(λ) represents the loss function based on either *L*_∞_ or *L_F_* norm which are defined by

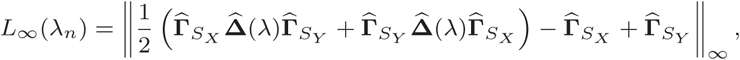

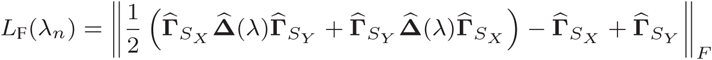

### 2.1 Simulation studies

In this section, we introduce simulation designs for evaluating the performance of the proposed matrix-variate differential network estimation method. For the temporal covariance matrices, we considered the following two types of structures. Type 1 is an autoregressive model,**Σ**_*T_X_*_ = (σ_*T_X_*,*i,j*_) with elements σ_*T_X_*,*i,j*_ = 0.4^|*i−j*|^ and **Σ**_*T_Y_*_ = (σ_*T_Y_*, *i,j*_) with elements σ_*T_Y_*, *i,j*_ = 0.5^|*i−j*|^, 1 ≤ *i,j* ≤ *q*. Type 2 is a moving average model, **Σ**_*T_X_*_ = (σ_*T_X_*, *i,j*_) with nonzero elements σ_*T_X_*, *i,j*_ = 1/(|*i −j*|+1) for |*i −j*|≤ 3 and **Σ**_*T_Y_*_ = (σ_*T_Y_*, *i,j*_) with nonzero elements σ_*T_Y_*, *i,j*_ = 1/(|*i −j*|+1) for |*i −j*|≤4. Note that **Σ**_*T_X_*_ and **Σ**_*T_Y_*_ are both nuisance parameters. We mainly focus on the spatial graphs of two groups encoded in **Ω**_*S_X_*_ and **Ω**_*S_Y_*_ respectively. We first generate the edge set E_*S_X_*_ for the group ***X***. To this end, we consider the following three graph structures: a hub graph, a scale-free graph, and a small-world graph, as shown in Figure 1.

**Fig. 1.**
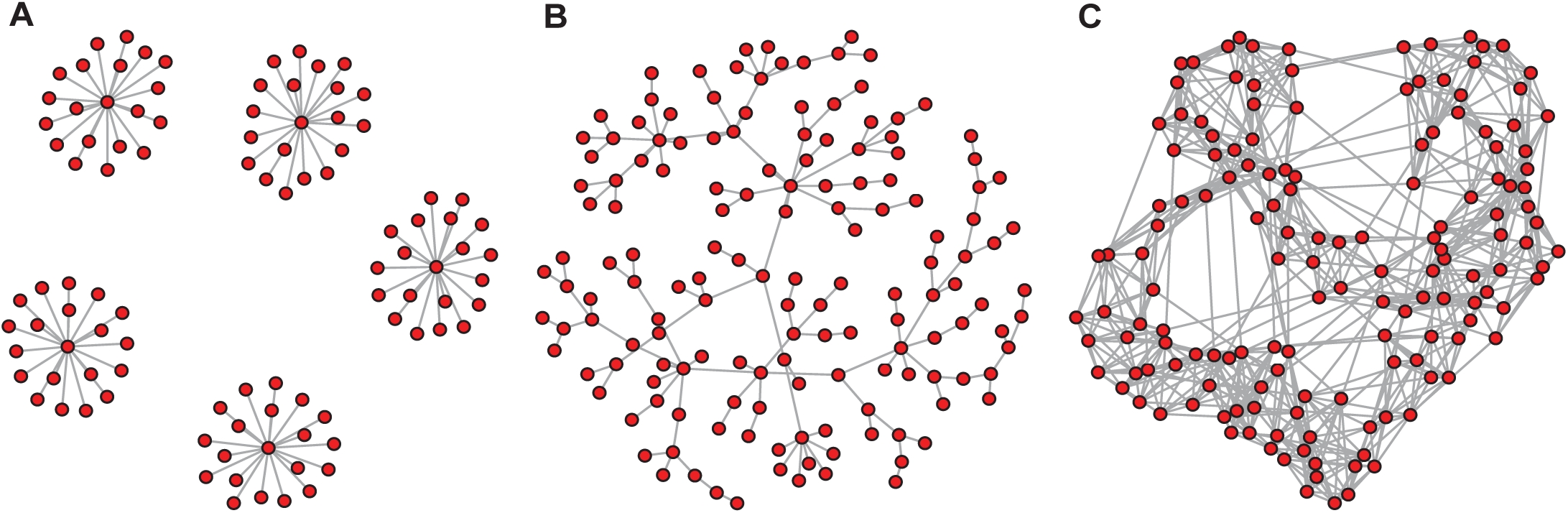
Three types of graph used in our simulation studies: (A) Hub graph; (B) Scale-free graph; (C) Small-world graph

- **Scenario 1: Hub graph** We partition *p* features into 5 equally-sized and non-overlapping sets: *C*_1_ ∪ *C*_2_ ··· ∪ *C*_5_ = {1,…,*p*}, |*C_k_*| = *p*/5, *C_i_* ⋂ *C_j_* = ∅. For the smallest *i* ∈ *C_k_*, we set (*i, j*) ∈ E_*S_X_*_ for all {*j ≠ i : j* ∈ *C_k_* }.
- Scenario 2: Scale-free graph Initially only one edge exists in E*_S_X__*. Then the graph grows such that each new node is connected to one existed node with the probability proportional to the degree of the node. In practice, we use R package “huge” to generate such a graph structure.
- Scenario 3: Small-world graph The graph is generated by the R package “rags2ridges” with 5 starting neighbors and 5% probability of rewiring (for further detail, one may refer to Wieringen and Peeters (2016)).

The non-zero entries of **Ω**_*S_X_*_ is then determined by the edge set E_*S_X_*_. The value of each nonzero entry of **Ω**_*S_X_*_ was generated from a uniform distribution with support [−0.5, −0.1]∪[0.1, 0.5]. To ensure positive definiteness of **Ω**^*S*^*X*, let **Ω**_*S_X_*_ = **Ω**_*S_X_*_ +(1+ |λ_min_(**Ω**_*S_X_*_)|)**I**. We then proceed to generate the differential matrix **Δ**_0_. For Scenario 1, we randomly select two hub nodes from the 5 equally-sized and non-overlapping sets and the differential matrix **Δ**_0_ is generated such that the connections of these two hub nodes change sign between **Ω**_*S_X_*_ and **Ω**_*S_Y_*_. For Scenario 2 and Scenario 3, we randomly select 40% of the edges in E_*S_X_*_ and the differential matrix **Δ**_0_ is generated such that the corresponding connections change sign between **Ω**_*S_X_*_ and **Ω**_*S_Y_*_. The covariance matrix **Σ**_*S_X_*_ and **Σ**_*S_Y_*_ are generated by (**Ω**_*S_X_*_)^−1^ and (**Ω**_*S_Y_*_)^−1^, respectively. Finally we generate *n*_1_ i.i.d observations of ***X*** from the *N_p × q_*(**0**, **Σ**_*T_X_*_ ⊗ **Σ**_*S_X_*_) distribution and *n*_2_ i.i.d observations of Y from the *N_p × q_*(**0**, **Σ**_*T_Y_*_ ⊗ **Σ**_*S_Y_*_) distribution.

In the simulation study, we examined a range of spatial and temporal dimensions and the sample sizes: *p* = {50, 100, 200}, *q* = {50, 100}and *n*_1_ = *n*_2_ = {20, 50}, which are consistent with the usual setup in functional connectivity analysis. All the simulation results are based on 100 replications.

We evaluate the performance of the estimation methods from the view of support recovery. The support recovery results are evaluated by true positive rate (TPR), true discovery rate TDR and true negative rate (TNR) along a range of tuning parameter λ. Suppose the true difference matrix **Δ** has the support 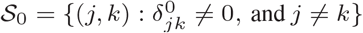 and its estimator 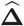 has the support set 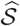. TPR, TDR and TNR are defined as follows:

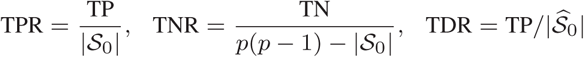

where TP and TN are the numbers of true positives and true negatives respectively, which are defined as

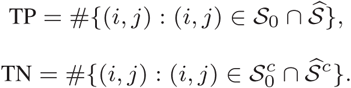

We compare the proposed Matrix-Variate Differential Model (denoted as MVDN) estimation method with the joint multiple matrix Gaussian graphs estimation method in Zhu and Li (2018). Note that in Zhu and Li (2018), non-convex and convex penalty are proposed to jointly estimate matrix graphs and we simply denote the corresponding method as Non-convex and Convex. Moreover, Zhu and Li (2018) shows the advantage of directly working with the matrix data rather than working with the vector-valued data after whitening in their simulation study, so there is no need to compare the matrix-variate methods with vector-valued methods such as Zhao *et al.* (2014), Cai *et al.* (2016) and He *et al.* (2017b).

### 2.2 Evaluation of computational complexity

We carried out simulations for Scenario 1 to compare the computing speed of different methods with *n* = {20, 50}, *q* = {50, 100} and *p* = {50, 100, 200, 300}. Timings (in seconds) are computed using a single core, Intel(R) Xeon(R) CPU E5-2697 v3 at 2.60GHz and 128 Gbytes memory, which is shown in Table 1.

**Table 1.** Average timings (in seconds) over 20 replications.

| $N = 20$ | | | | |
| --- | --- | --- | --- | --- |
| $p$ | $q = 50$ | | $q = 100$ | |
|  | MVDN | Non-convex | MVDN | Non-convex |
| 50 | 0.13 | 28.73 | 0.13 | 26.74 |
| 100 | 0.61 | 141.30 | 0.59 | 114.18 |
| 200 | 3.34 | 1079.23 | 2.98 | 676.28 |
| 300 | 10.27 | 4418.82 | 9.97 | 2262.24 |
| $N = 50$ | | | | |
| $p$ | $q = 50$ | | $q = 100$ | |
|  | MVDN | Non-convex | MVDN | Non-convex |
| 50 | 0.14 | 25.79 | 0.15 | 26.01 |
| 100 | 0.60 | 108.45 | 0.60 | 112.76 |
| 200 | 2.86 | 574.25 | 2.76 | 616.70 |
| 300 | 8.40 | 1624.75 | 7.87 | 1913.39 |

### 2.3 Application to Electroencephalography (EEG) data

The EEG data was collected in a study examining EEG correlates of genetic predisposition to alcoholism. It is available at website http://kdd.ics.uci.edu/datasets/eeg/eeg.data.html, which consists of 77 alcoholic individuals and 45 controls. Each subject was fitted with a 61-lead electrode cap and was recorded at 256 Hz for 1 second. There were in addition a ground and two bipolar deviation electrodes, which are excluded from the analysis. The electrode positions were located at standard sites (Standard Electrode Position Nomenclature, American Electroencephalographic Association 1990), and were organized into frontal, central, parietal, occipital, left temporal, and right temporal regions (Acharya *et al.* (2016)). Each subject performed 120 trials under three types of stimuli. For more details on data collection, one may refer to Zhang *et al.* (1995). Similar to Xia and Li (2017), we first preprocessed the data by first averaging all trials under a single stimulus condition following Li *et al.* (2010) and then performed an α-band filtering on the signals following Hayden *et al.* (2006). Finally, we obtain 61 × 256 matrix data for each subject.

## 3 Results

### 3.1 Theoretical results

We show that the proposed estimator 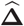 in (1) shares exactly the same support as the true differential partial correlation matrix **Δ** with probability tending to 1 under mild and regular conditions, indicating that the proposed method can identify all differential edges accurately as dimensions of matrix-valued data *p, q* and sample size *n* go to infinity. In fact, we show that as long as the signal strength is larger than 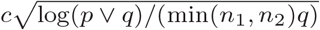 for some constant *c*, then we establish a theorem that 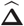 can recover not only the support of **Δ** but also the signs of its nonzero entries.

### 3.2 Simulation Results

In this section we present the simulation results for the experimental design introduced in Section 2.1. To evaluate the support recovery performance of the methods, we plot the ROC curve and the Precision-Recall (PR) curve for various scenarios. Figure 2 depicts the ROC curve and PR curve for Scenarios 1 (Hub Graph) with *p* =50, 100, 300 and autoregressive temporal covariance structure, in which the solid line represents MVDN, dotted line and dashed line represents Convex method and Non-convex method in Zhu and Li (2018) respectively. The top panels in Figure 2 are the ROC curves for *p* =50, 100, 200, from which we can see that the MVDN performs uniformly better than the Convex and Non-convex methods. It also can be seen that the Convex and Non-convex methods performs nearly the same and both worse than the MVDN method. The advantage of the MVDN method is obviously shown from the Precision-Recall curves in the bottom panels of Figure 2. When the true positive rate (TPR) does not exceed 80%, the true discovery rate (TDR) of MVDN method is always 100% while that of Convex and Non-convex methods do not exceed 50%. From Figure 2, we can also see that as dimension *p*, increases, the performances of all methods get worse, which is expected as the graph becomes large. What we want to emphasize is that the MVDN method generally performs robustly as *p* varies. Similar conclusions above from Figure 2 can be drawn from Figure 3 and Figure 4, which correspond to Scenario 2 (Scale-free graph) and Scenario 3 (Small-world graph) respectively. From Figure 2 to Figure 4, we can see that all methods performs similarly for different types of graph structures. To save space, we put all the other simulation results in the Supplementary material. From the Figures in the Supplementary material, we can also draw conclusion that for the same Scenarios, as sample size *n*_1_, *n*_2_ increases (with *p,q* fixed), the performances of all methods get better. Besides, with *n, p* fixed, and *q* increases, the performances of all methods also get better as we collect more temporal data, thus have more information excavated.

**Fig. 2.**
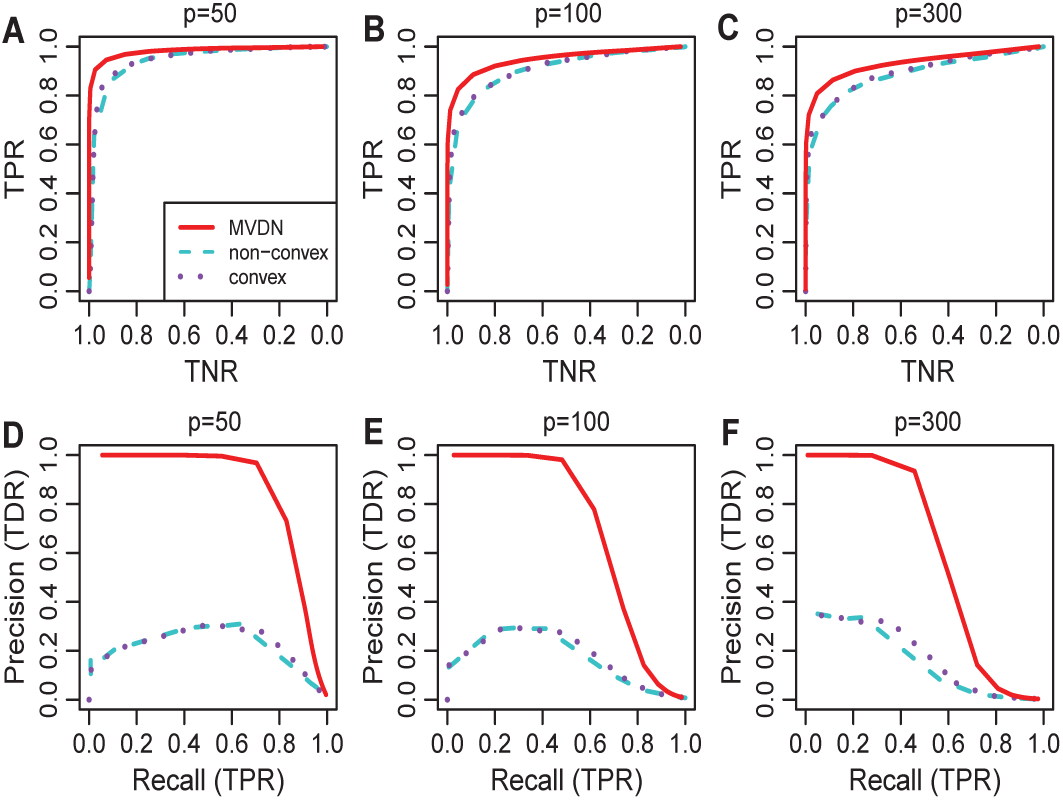
ROC curve (upper panels) and Precision-Recall curve (lower panels) for Scenario 1 with *p* =50, 100, 300 and autoregressive temporal covariance structure. Solid line represents MVDN, dotted line represents convex method, and dashed line represents non-convex method. The sample size *n*_1_ = *n*_2_ = 20 and *q* = 50.

**Fig. 3.**
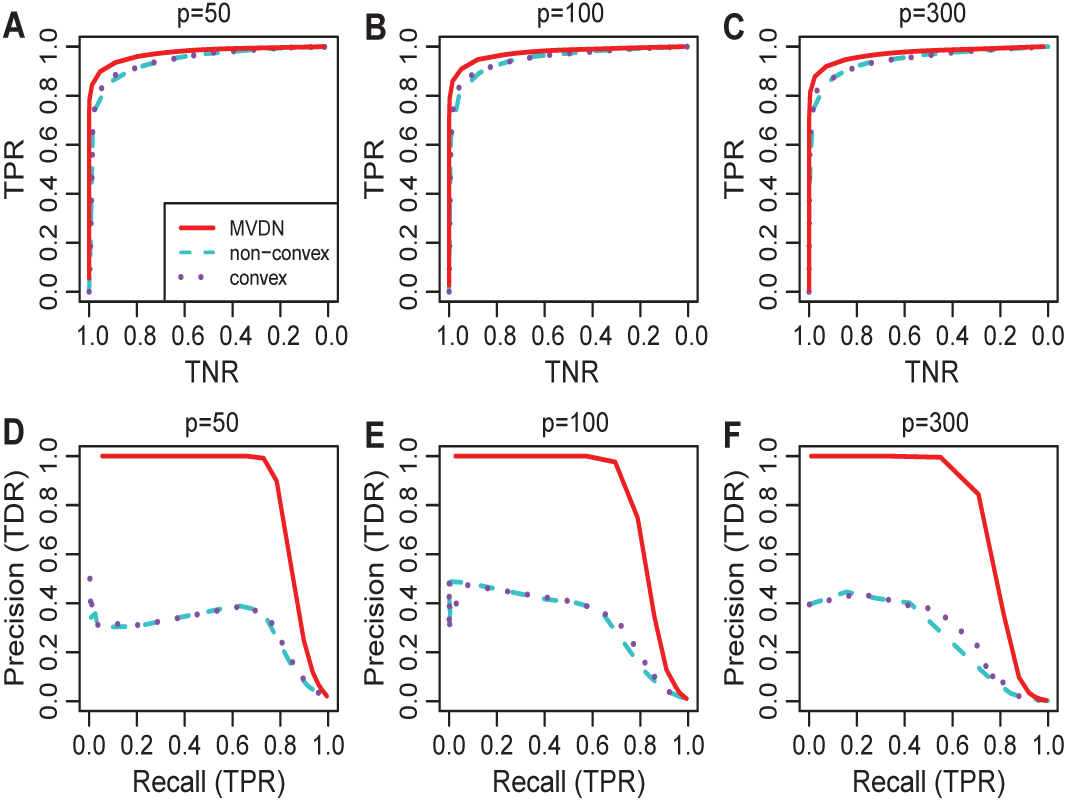
ROC curve (upper panels) and Precision-Recall curve (lower panels) for Scenario 2 with *p* = 50, 100, 300 and autoregressive temporal covariance structure. Solid line represents MVDN, dotted line represents convex method, and dashed line represents nonconvex method. The sample size *n*_1_ = *n*_2_ = 20 and *q* = 50.

**Fig. 4.**
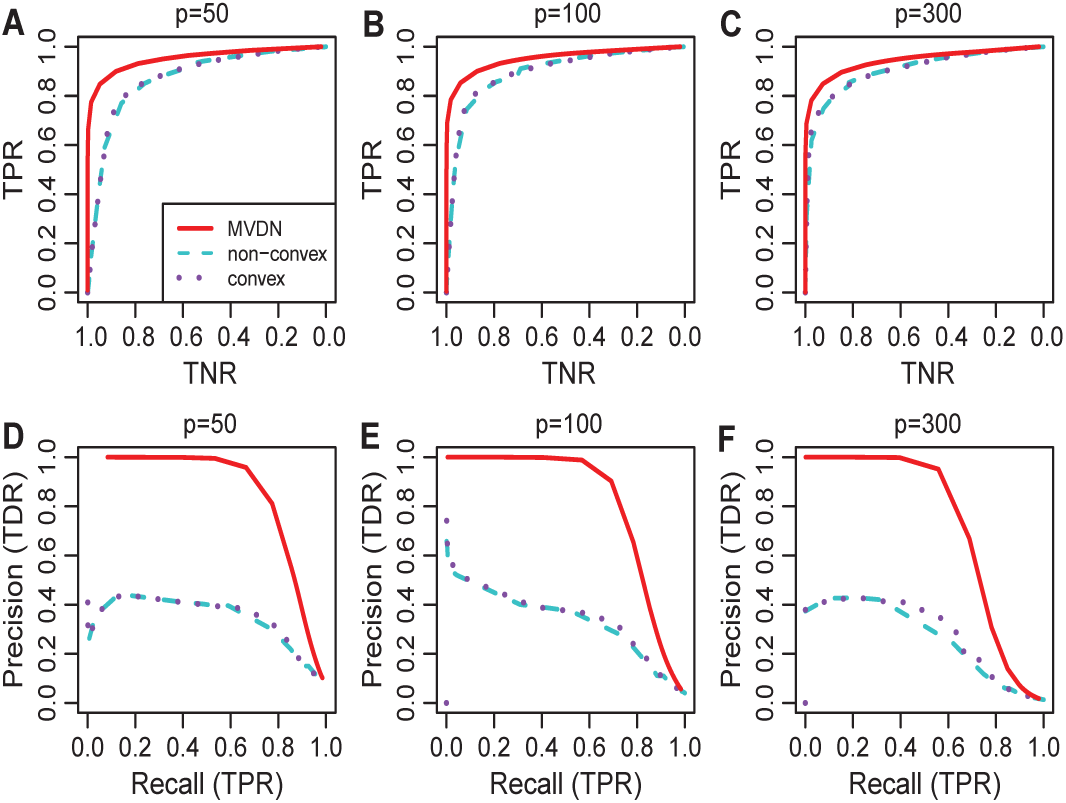
ROC curve (upper panels) and Precision-Recall curve (lower panels) for for Scenario 3 with *p* =50, 100, 300 and autoregressive temporal covariance structure. Solid line represents MVDN, dotted line represents convex method, and dashed line represents non-convex method. The sample size *n*_1_ = *n*_2_ = 20 and *q* = 50.

In summary, the simulation results show that the MVDN method always outperforms its competitors Non-convex and Convex method in all the scenarios, illustrating the advantages of the MVDN method. Zhu and Li (2018) showed that the non-convex and convex methods substantially outperforms the state of the art vector-normal-based methods. It further indicates the advantage of MVDN method against the vector-normal-based methods.

### 3.3 Application Results

Figure 5 illustrate the differential network estimated by different methods, in which orange edges show an increase in partial correlation dependency from control group to alcoholic group; grey edges show a decrease and red points stand for hub nodes whose number of edges ranks the top 5 for each method. Specifically, the tuning parameter λ for method MVDN is selected by minimizing the Bayesian information criterion, i.e., λ is chosen to minimize (*n*_1_ + *n*_2_)*L*(λ)+log(*n*_1_ + *n*_2_)‖**Δ**‖_0_ where *L*(λ) represents the loss function based on LF norm introduced in section 2. For the Non-convex and Convex method, the λ is selected by minimizing a prediction criterion by using fivefold cross-validation as described in Zhu and Li (2018), i.e, first divide the data set into five parts *D*_1_,…,*D*_5_ for the control (denoted as group 1) and the alcoholic group (denoted as group 2). For each group *k,k* = 1,2, define 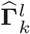 and 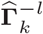 to be the sample correlation matrices based on samples in *D*_*l*_ and {*D*_1_,…,*D*_5_}\*D*_*l*_,*l* = 1,…,5. Let 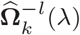 be the partial correlation matrices calculated based on 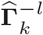 under the tuning parameter λ. Then λ is chosen to minimize *CV*(λ) which is defines as

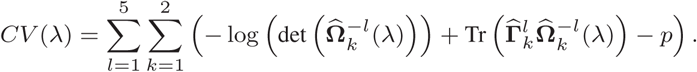

**Fig. 5.**
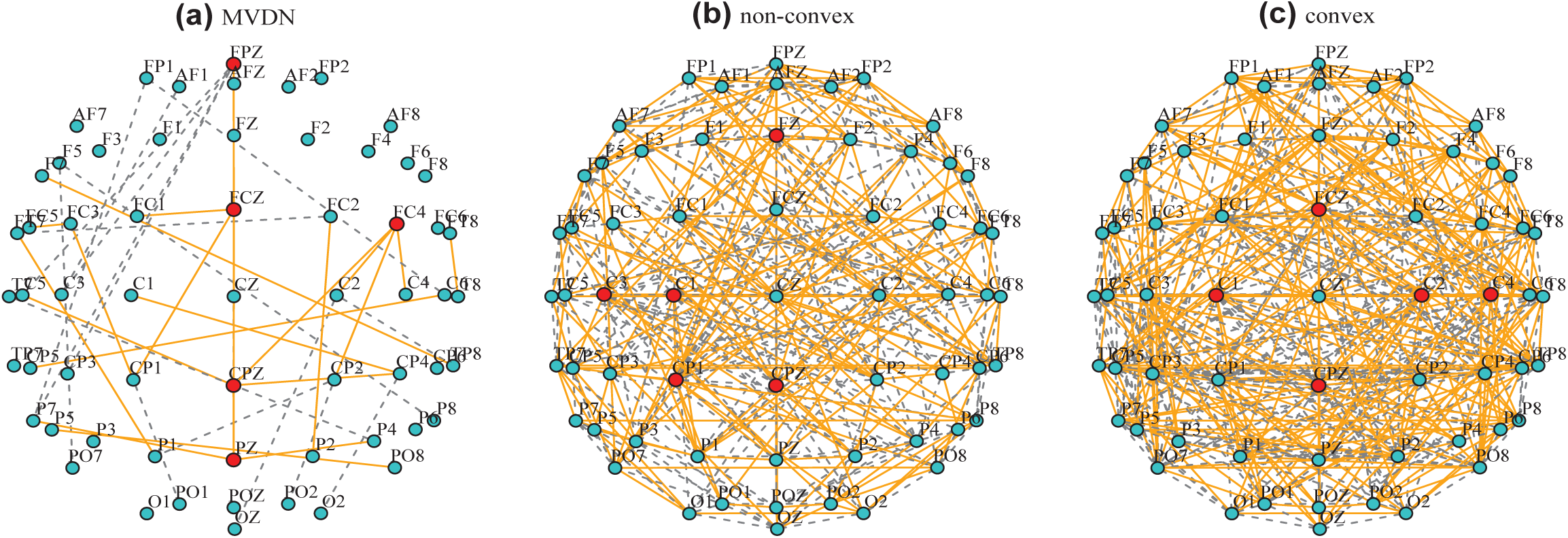
Differential network estimated by different methods. Orange edges show an increase in partial correlation dependency from control group to alcoholic group; grey edges show a decrease. Red points stand for hub nodes whose number of edges ranks the top 5 for each method.

From Figure 5, we can see that the differential network structure estimated by MVDN method is much sparse compared with those estimated by Non-convex and Convex methods. The differential network structure estimated by Non-convex and Convex methods are so dense that it is impossible to draw any useful conclusions from the graphs. In contrast, we can draw interesting conclusions from the differential graph estimated by MVDN. Firstly, we can see that those electrodes in the frontal region and temporal region (denoted by symbols FP, AF, F, TC and TP) show a decrease in connections and some asymmetry between the left and right frontal regions in the alcoholic group compared to the control, which agrees with the findings in the literature, see for example, Hayden *et al.* (2006). Secondly, those electrodes in the central region and parietal region mainly show a increase in connections in the alcoholic group compared to the control. Thirdly, about 56.1% of the identified differential edges by MVDN show a increase in connection from the control group to the alcoholic group, which indicates that alcoholic would strengthen the connection activities in brain region.

## 4 Discussion

Brain functional connectivity reveals the synchronization of brain systems through correlations in neurophysiological measures of brain activity. Growing evidence shows that the brain connectivity network experiences alternations with the presence of numerous neurological disorders. Thus it entails the analysis of differential network. The most typical feature of brain connectivity data is its matrix-form while the primary interest is to infer the spatial connectivity network. To our knowledge, no literatures focus on the differential network analysis for matrix-form type data. This motivates us to consider such an important topic. In this paper, we establish a matrix-variate differential network Model assuming that the matrix-variate data follows a matrix-normal distribution, and propose to exploit the D-trace loss function and a Lasso-type penalty to estimate the spatial differential partial correlation matrix directly. Theoretically we show that the proposed method can identify all differential edges accurately with probability tending to 1 in high-dimensional setting under mild and regular conditions.

The work could be extended in the following aspects. First, in this paper, we exploit the D-trace loss function and a Lasso-type penalty to estimate the spatial differential partial correlation matrix. In fact we can consider the constrained *l*_1_ minimization approach as in Zhao *et al.* (2014), that is,

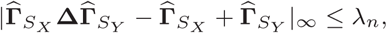

where λ_*n*_ is a tuning parameter. The theoretical properties of the estimator are still needed to be investigated, but we expect that the empirical performance should be similar.

Second, the matrix-normal distribution assumption for the brain connectivity data could be somewhat restrictive. Ning and Liu (2013) propose a semiparametric extension of the matrix-normal distribution, called the matrix-nonparanormal distribution. Thus we can assume that the brain connectivity data follows matrix-nonparanormal distribution and establish the differential network model for matrix-nonparanormal data. This extension is more challenging because there exists diverging number of unknown transformation functions needed to be carefully dealt. We leave this problem as a promising future work as there is a lot of work still needed to be done.

The proposed matrix-variate differential network model is very flexible and may provide deeper understanding of the brain connectivity alternation mechanism. It is demonstrated the matrix-variate differential network models enjoy great advantages over existing models and thus are highly recommended in the brain connectivity analysis.

## Acknowledgements

We would like to thank professor Yunzhang Zhu for providing the simulation codes of their Non-convex and Convex method.

